# The Social Cognition Paradox in Long-Duration Spaceflight: A VEN Fatigue Hypothesis for Duration-Dependent Emotion Recognition Decline

**DOI:** 10.64898/2026.08.06.743343

**Authors:** Esila Keskin, Margaret Windy McNerney, Nilufar Ali

## Abstract

Long-duration spaceflight may alter social cognition, yet the underlying biological mechanisms remain unclear. Emotion Recognition Task (ERT) performance remains stable during typical 6-month International Space Station (ISS) missions but declines markedly during the 340-day NASA Twins Study, suggesting duration-dependent vulnerability. Here, we propose the Von Economo Neuron (VEN) Fatigue Hypothesis, which posits that microgravity increases demand on VEN-associated social cognitive networks, eliciting adaptive myelination during shorter missions before compensatory mechanisms fail with prolonged exposure. To evaluate this hypothesis, we integrated evidence from rodent, human cortical organoid, astronaut plasma proteomic, and neuroimaging datasets. ISS-flown rodent frontal cortex demonstrated increased expression of myelination-related genes, while spaceflown cortical organoids exhibited changes consistent with oligodendrocyte remodeling. Astronaut plasma transcriptomics identified reproducible alterations in VEN-associated and myelination-related proteins across independent missions, and resting-state fMRI revealed transient changes in frontal insula connectivity following long-duration spaceflight. Together, these findings provide convergent evidence supporting the VEN Fatigue Hypothesis and identify adaptive myelination and VEN-associated network remodeling as candidate mechanisms underlying duration-dependent changes in social cognition during long-duration spaceflight.

## 1. Introduction

A striking paradox defines spaceflight social cognition. In the single-subject 340-day NASA Twins Study (Garrett-Bakelman et al., 2019), Emotion Recognition Task (ERT) speed, a validated measure of social cognitive processing speed (Basner et al., 2015), shows the largest early-to-late inflight change of any cognitive speed domain (−1.8 SD), while spatial orientation and visual memory remain stable or improve at the same mission phase. Yet in 6-month ISS missions (N = 25), ERT speed shows effectively no inflight change (+0.106 SD) (Dev et al., 2024). General cognitive fatigue, radiation, or confinement stress would be expected to affect multiple domains simultaneously; the concurrent stability of spatial and memory domains in both datasets cannot be explained by a non-specific account (Basner et al., 2015; Strangman et al., 2014). No mechanistic explanation of the ERT-specific duration-dependent pattern has been proposed. We treat the single-subject Twins Study finding throughout as a hypothesis-generating case observation, corroborated by the stable 6-month group trajectory, and not as a basis for population-level inference. ERT specifically engages the anterior cingulate cortex (ACC) and frontal insula (Allman et al., 2011), the only brain regions housing von Economo neurons (VENs). VENs are large bipolar projection neurons (soma 65-80 µm) with a sparse afferent fan-in of approximately 10-100 synapses versus ~10,000 for standard pyramidal neurons, and thick myelinated axons (Allman et al., 2011; Butti et al., 2013; Figure 1). This morphology is optimized for fast sparse signal transmission. Because their afferent pool is so small, degraded or novel social inputs that would be averaged out across thousands of pyramidal synapses instead drive strong VEN responses, making VENs exquisitely sensitive to input quality. VENs are selectively depleted in frontotemporal dementia, producing profound social intuition loss, yet preserved in Alzheimer’s disease, which spares social cognition (Seeley et al., 2006; Stimpson et al., 2011). The Fast Lane Hypothesis (Keskin, 2026) formalises this circuit logic, proposing that VENs implement a fast pathway for social decisions whose integrity specifically governs social processing speed rather than representational capacity. VENs are phylogenetically restricted to species with large, complex social groups. They are absent in rodents, sparse in great apes, and most numerous in humans, with ACC VEN density correlating positively with social group size across primate species (Allman et al., 2011; Stimpson et al., 2011). This evolutionary distribution implies VENs specifically support cognitive demands that scale with social complexity, precisely the demands most radically altered by the gravitational and social context disruption of long-duration spaceflight.

**Figure 1.**
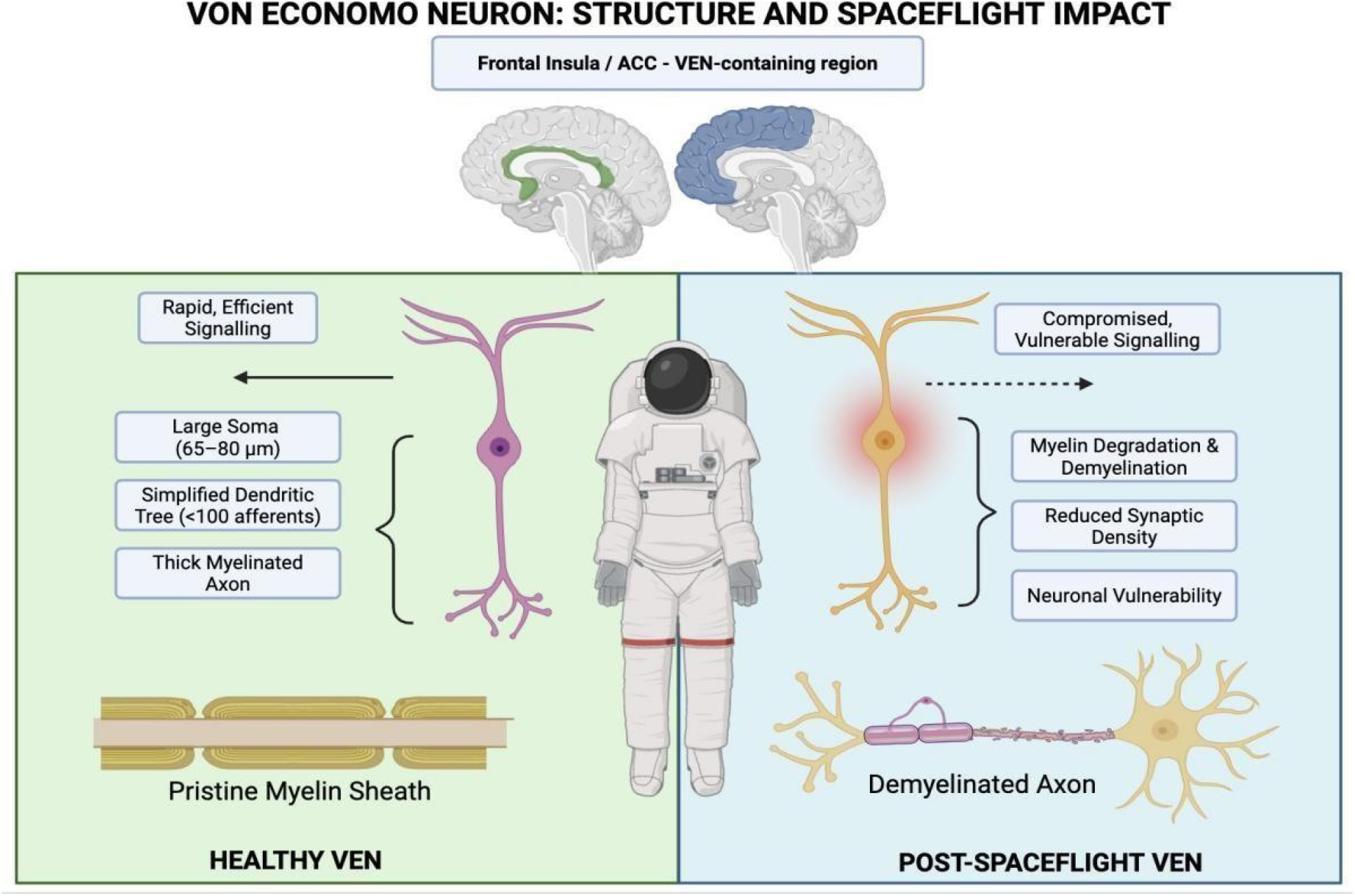
Von Economo neuron (VEN): structure and spaceflight impact. Left (Healthy VEN): Large bipolar soma (65-80 µm), simplified dendritic tree (<100 afferents), and thick myelinated axon enabling rapid, efficient signaling exclusively in the frontal insula and ACC (brain sections, inset: green = ACC; blue = frontal insula). Right (Post-Spaceflight VEN): The same cell class showing myelin degradation and demyelination, reduced synaptic density, and enhanced neuronal vulnerability, producing compromised signaling. Demyelinated axon (lower right) contrasts with pristine myelin sheath (lower left). The right panel depicts the proposed long-duration failure state when compensatory capacity is exceeded; the compensatory myelination upregulation phase, the primary molecular finding, is shown in Figures 4 and 5. Created in BioRender. Keskin, E. (2026) https://BioRender.com/gajlo2i.

We extend the Fast Lane Hypothesis to long-duration spaceflight. In microgravity, gravity-referenced cues that calibrate social gesture kinematics, eye contact geometry, and body orientation are fundamentally disrupted (Garrett-Bakelman et al., 2019). We propose that these degraded inputs fall outside the VEN calibration regime, driving disproportionate and sustained VEN firing. We term this the VEN Fatigue Hypothesis and derive two testable predictions.

### Hypothesis 1 (Molecular)

VEN myelination genes should be specifically upregulated in ISS frontal cortex as a compensatory response to sustained conduction demand, absent in ground-based analogues that replicate physical stressors but not gravity-referenced social cue disruption.

### Hypothesis 2 (Cognitive)

ERT speed should be stable within missions where myelination compensation is sufficient (≤6 months) and should decline specifically during longer missions where fatigue accumulates beyond compensatory capacity.

A structural prediction follows for organoid models: while some assembloid systems develop network activity, cortical organoids at this stage (approximately 30 days maturation) lack established mature VEN circuits and sustained social-signal-driven activity. They should therefore show direct microgravity effects on OPC differentiation rather than activity-driven compensatory myelination, potentially diverging in myelination direction from in vivo tissue (Espinosa-Jeffrey et al., 2013). We test Hypotheses 1 and 2 across five independent molecular datasets and additionally report resting-state fMRI evidence in human cosmonauts (Jillings et al., 2023) consistent with frontal insula circuit-level perturbation.

## 2. Methods

### 2.1 Data Sources

Seven datasets were analysed (Table 1); full protocols are in Supplementary Section S1. ISS rodent transcriptomics (GSE239336): GeoMx Digital Spatial Profiling of frontal cortex from rodents flown 35 days on the ISS (SpaceX CRS-24; spaceflight vs. ground control). Ground-based analogue (OSD-202; NASA Open Science Data Repository (OSDR); (Mao, 2018)): hindlimb unloading with cobalt-57 radiation vs. normal loaded control (1 month). Human iPSC-derived cortical organoids (GSE259421; Marotta et al., 2024): bulk RNA-seq from organoids maintained 38 days on ISS or in ground control (nLEO = 9, ngnd = 10 combined). Organoids at this stage (approximately 30 days of maturation) contain primarily OPCs and immature neurons rather than mature myelinating oligodendrocytes. Raw counts were normalised to CPM and log2-transformed; Ensembl identifiers were mapped to HGNC symbols via MyGene.info (Xin et al., 2016). Cortical organoid proteomics (PXD069807; Martins et al., 2026): Orbitrap Astral LC-MS/MS from iPSC brain organoids maintained 30 days on ISS, processed at 1% protein FDR. Cognitive data: Twins Study values digitised from Figure 10B of Garrett-Bakelman et al. (2019) (estimated uncertainty ±0.2 SD); 6-month data from Table 2 of Dev et al. (2024) (N = 25). Human blood transcriptomics: SOMA atlas (Overbey et al., 2024; <u>soma.weill.cornell.edu</u>), aggregating NASA Twins Study (year-long ISS) and Inspiration4 PBMC and cell-free RNA. Cosmonaut rsfMRI (Jillings et al., 2023; NeuroVault:12152; Gorgolewski et al., 2015): resting-state fMRI ICC from N = 15 cosmonauts before, shortly after, and 8 months after missions exceeding 3 months.

**Table 1.** Molecular, neuroimaging, and cognitive datasets analysed in this study.

| Dataset | Species / Cell Type | N | Assay | Comparison |
| --- | --- | --- | --- | --- |
| GSE239336 | Rodent frontal cortex | N=3 | GeoMx DSP spatial transcriptomics | Spaceflight vs ground control |
| OSD-202 | Mouse brain | N=6 | Bulk transcriptomics | Ground analogue vs normal control |
| GSE259421 | Human iPSC cortical organoids | N LEO=9, n gnd=10 | Bulk RNA-seq | 38-day ISS vs ground |
| PXD069807 | Human iPSC brain organoids | N = 2 (ISS) vs. N = 3 (Ground) | LC-MS/MS proteomics | ISS vs ground |
| SOMA atlas | Human astronaut blood | N = 1 (Twins Study, longitudinal) + N = 4 (Inspiration4 crew) | PBMC + CPT RNA | Inflight vs pre-flight |
| NeuroVault:12152 | Human cosmonauts | N=15 | Resting-state fMRI | Post-flight vs pre-flight, 8-month follow-up |
| NASA Twins Study | Single human subject | N=1 | Cognitive battery | Early vs late inflight (340 days) |

**Table 2.** SOMA atlas: VEN panel genes in human astronaut blood. Best (lowest p) comparison per gene across NASA Twins Study (year-long ISS; CPT) and Inspiration4 (I4). All eight genes reached p < 0.05.

| Gene | Category | Dataset | Comparison | $\log_2\text{FC}$ | $p$ | |
| --- | --- | --- | --- | --- | --- | --- |
| *MOG* | Myelination | Twin (CPT) | Inflight 2<sup>nd</sup> half vs ground | +5.82 | $4.59 \times 10^{-27}$ | * * * |
| *MBP* | Myelination | I4 PBMC | Post vs pre-flight | +0.60 | $\approx 0$ | * * * |
| *NEFL* | Fast Signalling | Twin (CPT) | Inflight 2<sup>nd</sup> half vs ground | +3.13 | $6.42 \times 10^{-5}$ | * * * |
| *VDAC1* | Metabolic Supp. | Twin (CPT) | Inflight 2<sup>nd</sup> half vs ground | -0.58 | $1.32 \times 10^{-4}$ | * * * |
| *SYP* | Metabolic Supp. | Twin (CPT) | Inflight 1<sup>st</sup> half vs ground | +0.73 | $1.29 \times 10^{-3}$ | ** |
| *SNAP25* | Metabolic Supp. | Twin (CPT) | Inflight 2<sup>nd</sup> half vs ground | +3.33 | $7.57 \times 10^{-3}$ | ** |
| *NEFH* | Fast Signalling | I4 cfRNA | Post vs pre-flight | +1.00 | $3.61 \times 10^{-2}$ | * |
| *NEFM* | Fast Signalling | Twin (CPT) | Inflight 2<sup>nd</sup> half vs ground | +1.03 | $4.27 \times 10^{-2}$ | * |

### 2.2 VEN Gene Panel

The 31-gene panel was defined a priori and prospectively deposited in the <u>Brain AWG GitHub repository</u> before any data access (April 2026; see also Keskin, 2026), across five categories: Myelination (*MBP, MOG, PLP1, MAG, CNP, MOBP, ERMN*); Fast Signalling (*SCN1A, KCNQ2, ANK3, NEFH, NEFM, NEFL, SNCG*); Social Circuit (*OXTR, AVPR1A, HTR2A, DRD1, CHRM1, GABRB2*); Layer V Projection (*FEZF2, BCL11B, TBR1, SATB2, CUX1*); Metabolic Support (*VDAC1, ATP2B2, SLC17A7, SNAP25, SYP, NRXN1*). Panel genes incorporated established VEN-specific markers identified from microdissected-cell RNA sequencing of human anterior cingulate cortex (Yang et al., 2019). No gene is exclusively expressed by VENs; signals reflect VEN-associated transcriptional programs.

### 2.3 Statistical Analysis

For GSE239336 and OSD-202, published log2FC values were used directly. For GSE259421, per-gene mean differences (ISS minus ground) were computed from log2(CPM + 1) without individual-gene filtering. Category-level permutation tests (N = 10,000 random gene sets of equal size; seed 42) assessed pathway specificity against the genome-wide background; results are reported as standard deviations above the null with two-tailed p-values. The permutation test assesses pathway specificity; the within-panel one-sample t-test assesses intra-category consistency; see Supplementary Section S2 for interpretation of cases where they diverge. The cross-domain cognitive comparison (Section 3.1) is a purely descriptive within-person extremity statistic; no population-level inference from a single-subject observation is intended.

## 3. Results

### 3.1 ERT Shows a Duration-Dependent Pattern: A Single-Subject Case Observation

In the 340-day NASA Twins Study (single subject), ERT speed showed the largest early-to-late inflight change of any of the ten NASA Cognition battery domains: −1.8 SD. Using the nine concurrent domain change scores as a within-person reference, ERT stood 11.41 standard errors below the person-level mean of concurrent domains (descriptive cross-domain extremity statistic; df = 8; no population inference intended; Figure 3). Spatial orientation (LOT: 0.0 SD) and visual memory speed (VOLT: +0.6 SD) were stable or improving at the same mission phase (Figure 2). Manual digitisation uncertainty (±0.2 SD) does not alter the conclusion: at the conservative lower bound (−1.6 SD), ERT remains the most extreme domain by more than 0.7 SD relative to the next largest change. In contrast, 6-month ISS missions (N = 25) showed ERT change of +0.106 SD which is effectively zero (Dev et al., 2024). The duration difference (340-day vs. 6-month) is −1.91 SD. We treat these findings solely as a hypothesis-generating case observation motivating the VEN Fatigue Hypothesis.

**Figure 2.**
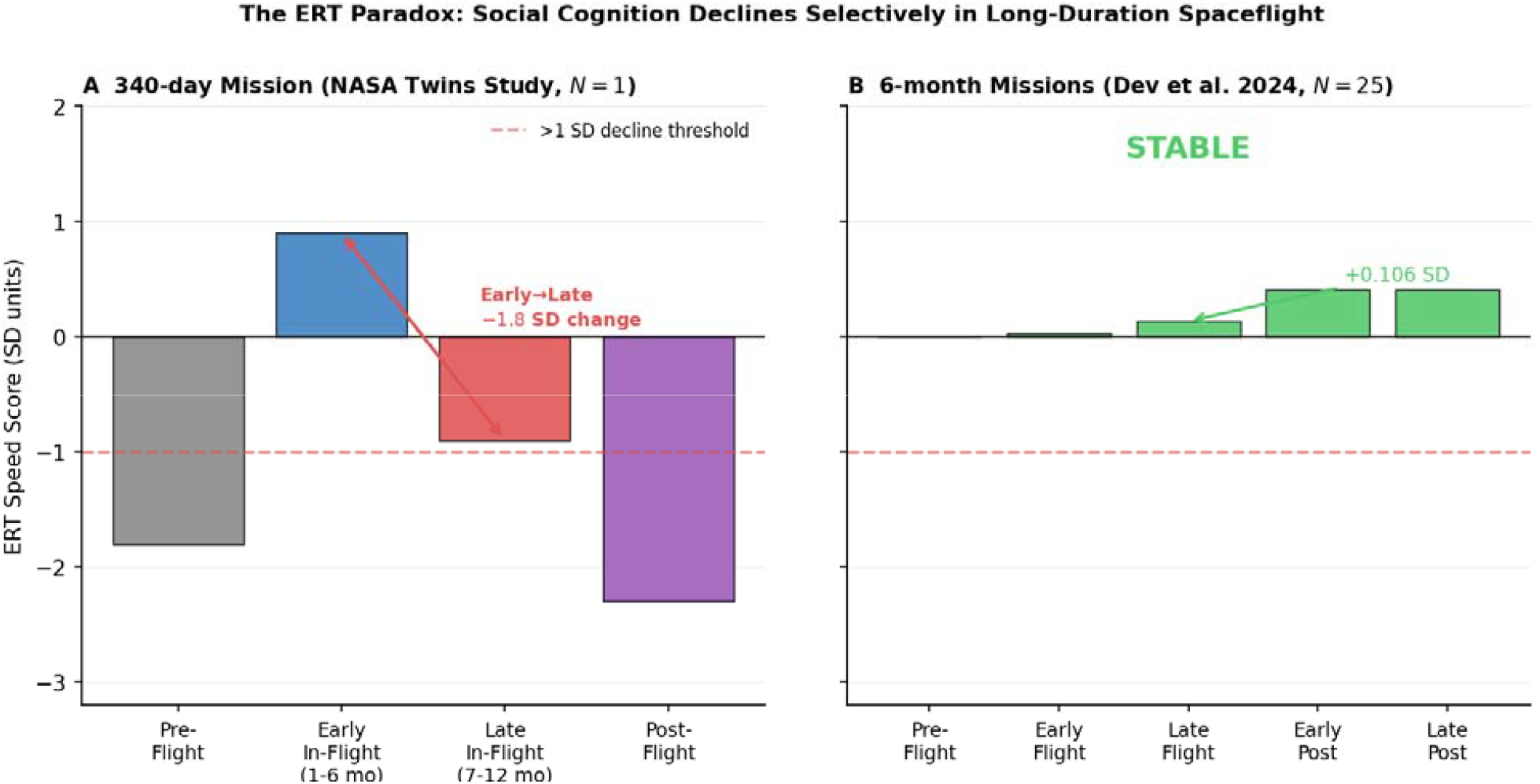
ERT shows a duration-dependent pattern (case observation). (A) In the single-subject 340-da NASA Twins Study (astronaut age: 51 years), ERT speed shows the largest inflight change of any cognitive domain (−1.8 SD), while spatial (LOT: 0.0 SD) and visual memory (VOLT: +0.6 SD) domains remain stable or improve (case-study observation; no population inference). (B) In 6-month ISS missions (N = 25; astronaut age range: 33–61 years, mean 45.1 years; Dev et al., 2024), ERT speed shows no systematic late-inflight decline (+0.106 SD) (Dev et al., 2024). Data from Garrett-Bakelman et al. (2019) and Dev et al. (2024); redrawn by E.K. from published values.

**Figure 3.**
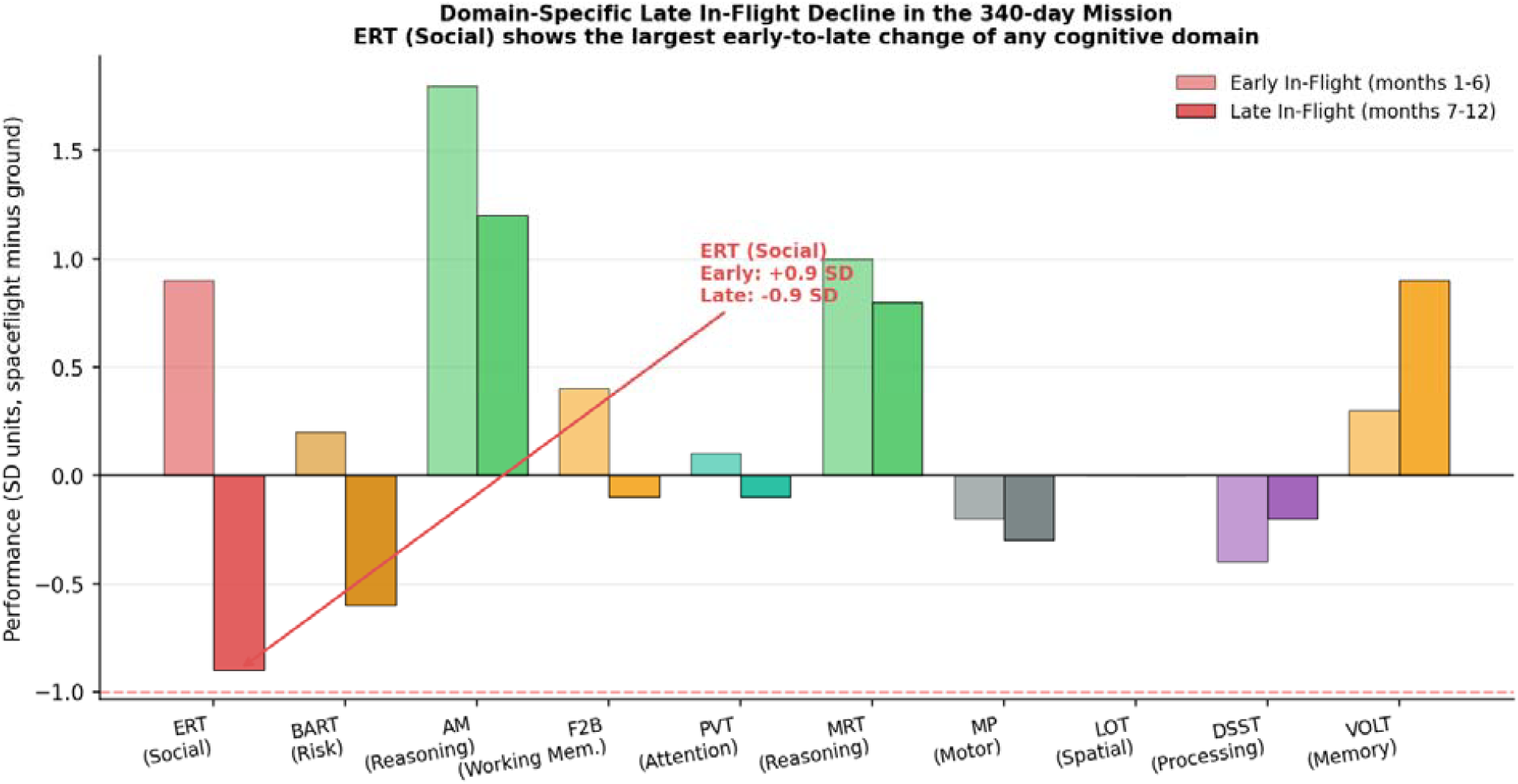
ERT is the most extreme domain across all ten cognitive speed tasks (single-subject cas observation). Early-to-late inflight change scores (SD units) for all 10 NASA Cognition battery subtests in the 340-day single-subject observation (Garrett-Bakelman et al., 2019). ERT (red) is the most extreme domain by >0.7 SD; spatial orientation (LOT: 0.0 SD) and visual memory (VOLT: +0.6 SD) are stable or improving at the same mission phase. Dashed line: within-person mean of the nine non-ERT domains. This is a single-subject case observation; no population inference is intended.

### 3.2 Spaceflight-Specific Myelination in ISS Rodent Frontal Cortex

In ISS frontal cortex (GSE239336), myelination genes showed a mean log2FC of +0.381 (t = 3.14, p = 0.025; n = 7 genes), 3.89 SD above the genome-wide permutation null (permutation p = 0.0001; Figure 4). Social circuit genes were also specifically upregulated (+0.227; 1.85 SD; p = 0.033). No other category reached permutation specificity. The permutation test drew 10,000 random gene sets of equal size from the full expressed transcriptome and compared their mean fold change to the panel mean, asking whether the myelination signal exceeded what chance sampling would predict. Spaceflight-induced transcriptomic changes in mouse brain have been documented in independent experiments (Kremsky et al., 2023), supporting the reproducibility of the ISS frontal cortex response.

**Figure 4.**
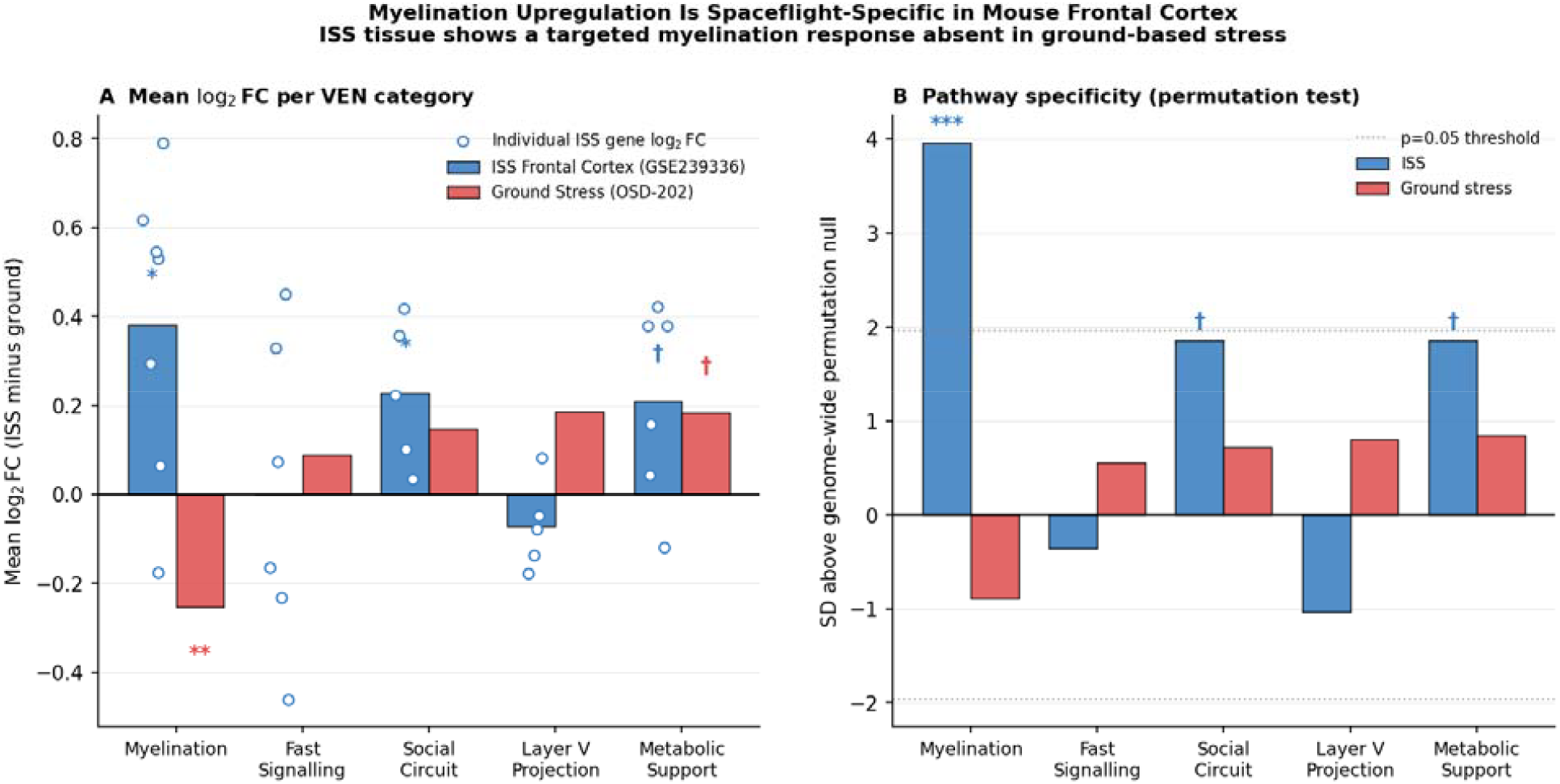
Myelination upregulation is spaceflight-specific in rodent frontal cortex. ISS tissue (GSE239336) shows a specific myelination signal 3.89 SD above the genome-wide permutation null (p = 0.0001). Ground-based analogue (OSD-202; hindlimb unloading + cobalt-57 radiation) shows no specific signal (−0.88 SD; p = 0.164), with directional reversal of the myelination mean. **(A)** Mean log2 fold change per VEN category (ISS minus ground); open circles show individual-gene log2FC values within the ISS panel, jittered horizontally for visibility. **(B)** The same comparison expressed as SD above the genome-wide permutation null, a category-level specificity statistic with no single-gene equivalent, hence no individual points; dotted lines mark ±1.96 SD, the two-tailed p=0.05 permutation threshold. ∗ ∗ ∗ p < 0.001; ∗ p < 0.05 (permutation, two-tailed, N = 10,000, seed 42).

In the ground-based analogue (OSD-202; hindlimb unloading with cobalt-57 radiation), the same panel showed mean log2FC of −0.254 which is not pathway-specific: −0.88 SD from the genome-wide null (permutation p = 0.164), with the panel following the global downregulation trend. The direction reverses between ISS (+0.381) and ground (−0.254), indicating qualitatively distinct rather than attenuated molecular responses. Because OSD-202 replicates radiation and mechanical unloading but not orbital microgravity’s disruption of gravity-referenced social and vestibular cues, this dissociation is consistent with Hypothesis 1 and with a gravity-dependent rather than radiation-dependent mechanism.

### 3.3 Human iPSC-Derived Cortical Organoids: Layer V Projection and Myelination Signals

All 31 panel genes and *NOS1* were detected in GSE259421 (full results in Supplementary Table S1). In the combined group (nLEO = 9, ngnd = 10): Layer V Projection genes showed the strongest and most specific signal: mean log2FC = +1.347 (6.17 SD above null; permutation p = 0.0008; Figure 5). Dominant contributors were *BCL11B* (+4.12) and *TBR1* (+2.52), master transcription factors for Layer V cortical identity (Molyneaux et al., 2007). High permutation significance with non-significant within-panel t-test (p = 0.189) reflects signal concentrated in two genes within a five-gene panel (see Supplementary Section S2). Metabolic Support genes were specifically upregulated (+0.731; 3.60 SD; p = 0.009), driven by *SLC17A7* (+2.13) and *NRXN1* (+0.88). Myelination genes were significantly downregulated (−0.456; −2.35 SD; p = 0.035), driven by *ERMN* (−1.60), *MBP* (−0.94), and *PLP1* (−0.85). This direction is opposite to the ISS rodent upregulation and constitutes the a priori predicted result: organoids at this stage contain primarily OPCs rather than mature myelinating oligodendrocytes, so their myelination signals reflect direct microgravity effects on OPC differentiation, not activity-driven compensatory upregulation. This prediction was registered before data access (<u>Brain AWG GitHub repository</u>, April 2026). Layer V and Metabolic Support signals replicated in dopaminergic organoids from the same dataset (7.12 SD, p = 0.0002; Supplementary Table S2 and Section S4), confirming the response is not restricted to the cortical lineage. *NOS1* (neuronal nitric oxide synthase), a direct biochemical VEN marker, showed +0.78 log2FC in dopaminergic organoids, consistent with dopaminergic stress responses documented in spaceflight (Ali et al., 2025, 2026). In the dopaminergic sub-dataset, the Layer V signal was numerically stronger than in cortical organoids, driven by the same *BCL11B* and *TBR1* transcription factors; myelination genes were similarly downregulated. That an identical transcriptional programme is activated across lineages that diverge substantially in mature identity indicates the spaceflight response targets progenitor-stage regulatory networks shared between VEN-adjacent neuronal classes, rather than a cortical-specific fate programme, and suggests the effect acts during an early developmental window common to both. Proteomics (PXD069807; Martins et al., 2026) confirmed VEN-associated structural and metabolic proteins are detectable in these organoids; all followed the global ISS downregulation trend with no VEN-pathway-specific signal, consistent with the Layer V transcription factors driving the organoid result falling below the mass-spectrometry detection threshold (Supplementary Section S5).

**Figure 5.**
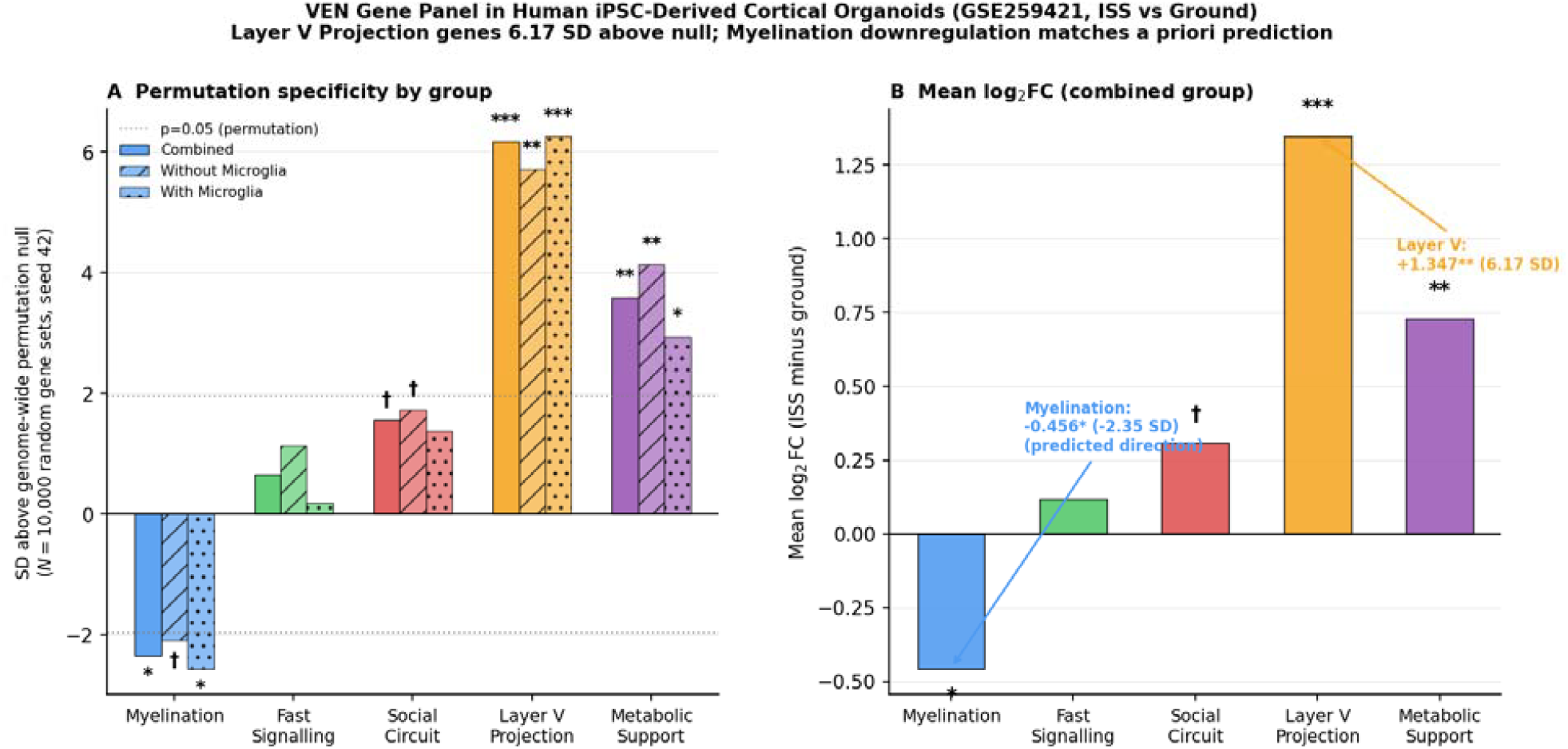
VEN gene panel permutation specificity in human iPSC-derived cortical organoids (GSE259421). Layer V Projection genes are the most strongly and specifically upregulated category (6.17 SD; p = 0.0008). Myelination genes are significantly below null (−2.35 SD; p = 0.035), the a priori predicted result for OPC-stage organoid models. **(A)** SD above the genome-wide permutation null for the combined organoid group and two microglia-inclusion sensitivity subsets (without microglia; with microglia); dotted lines mark ±1.96 SD, the two-tailed p=0.05 permutation threshold. **(B)** Mean log2 fold change (ISS minus ground) for the combined group. ∗ ∗ ∗ p < 0.001; ∗∗ p < 0.01; ∗p < 0.05; † p < 0.10 (N = 10,000 permutations, seed 42).

### 3.4 Human Evidence: Blood Transcriptomics and Cosmonaut Neuroimaging

SOMA atlas (human astronaut blood). All eight VEN panel genes queried from the SOMA atlas (Overbey et al., 2024) reached p < 0.05 in their best available comparison (Table 2; Figure 6). The myelination signal was unambiguous: *MOG* showed log2FC = +5.82 (p = 4.59×10^−27^) in year-long ISS CPT blood cells, replicating in Inspiration4 PBMC (*MBP*: log2FC = +0.60, p ≈ 0), two independent missions spanning different durations and cohorts. Fast Signalling genes (*NEFL*: +3.13; *NEFM*: +1.03; *NEFH*: +1.00) and Metabolic Support genes (*VDAC1*: −0.58; *SNAP25*: +3.33; *SYP*: +0.73) all reached significance. *MOG* expression in peripheral blood is near-zero at baseline, so the large log2FC carries absolute-scale caveats; cross-mission replication across independent cohorts and distinct sample types strengthens confidence. Myelination upregulation being detectable across both short-duration (Inspiration4, 3 days) and long-duration (year-long ISS) missions is consistent with Hypothesis 2: the blood signal reflects active compensation occurring throughout spaceflight, whereas selective ERT speed decline is predicted only when this compensation eventually fails at longer mission durations.

**Figure 6.**
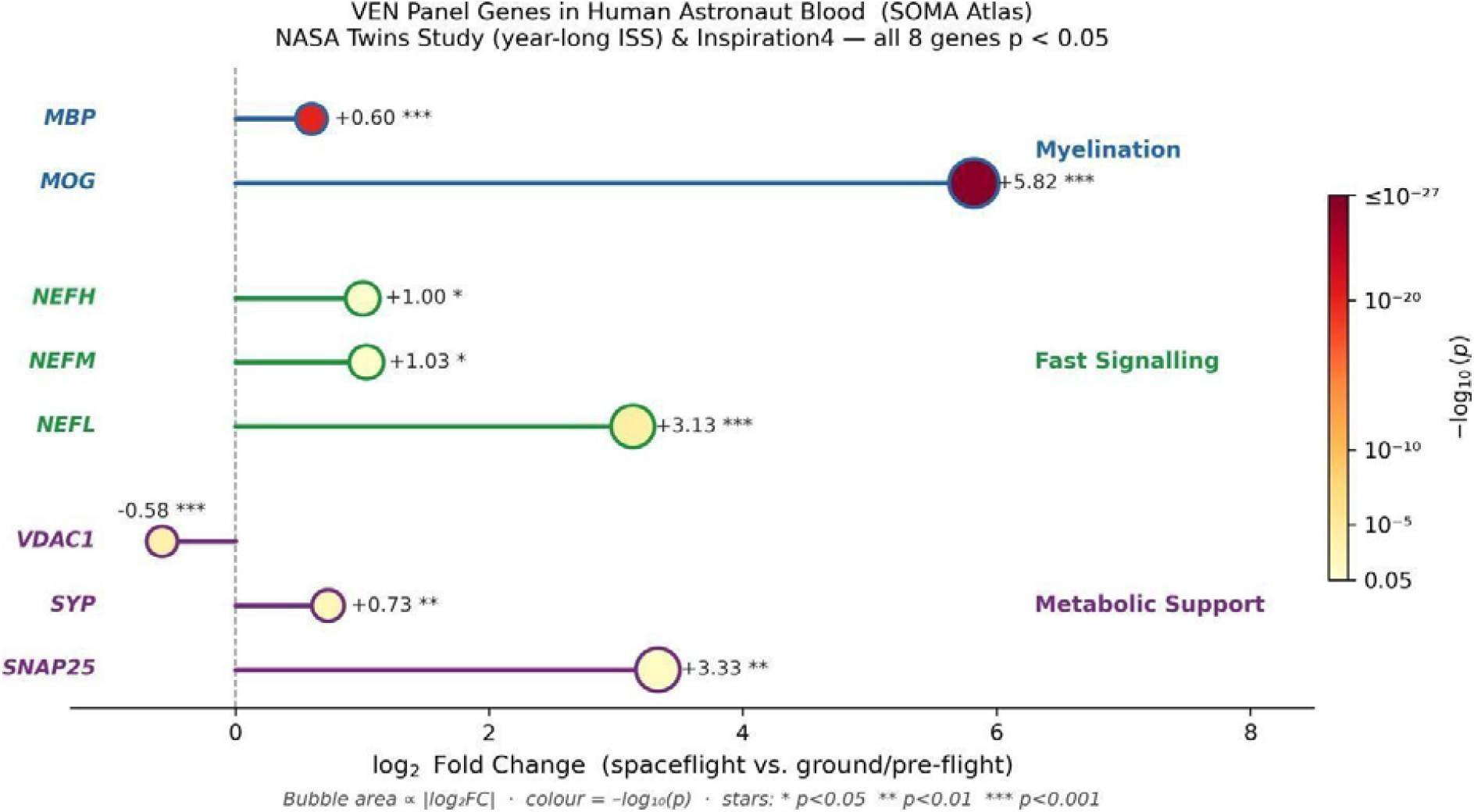
VEN panel genes in human astronaut blood (SOMA atlas). log2 fold change for eight VEN panel genes (Overbey et al., 2024; <u>soma.weill.cornell.edu</u>). All eight genes reached p < 0.05. Myelination genes (MOG, MBP) replicate across two independent missions (NASA Twins Study year-long ISS and Inspiration4). Significance: ∗ ∗ ∗ p < 0.001; ∗∗ p < 0.01; ∗ p < 0.05.

#### Cosmonaut resting-state fMRI

Jillings et al. (2023) acquired resting-state fMRI from cosmonauts (N = 15) before, shortly after, and 8 months after missions exceeding 3 months; of these, n = 11 completed the 8-month follow-up scan, and the normalisation finding is drawn from this subset. In the ICC map of connectivity changes that normalise to preflight levels by the 8-month follow-up, the two largest clusters localise bilaterally to the frontal insula and adjacent frontal operculum (right: 540 voxels, peak MNI (+48, +2, −6), t = −5.91; left: 428 voxels, peak MNI (−36, −16, 0), t = −5.91; NeuroVault:12152). The frontal insula is one of the two brain regions housing VENs. Insula connectivity was therefore specifically altered by spaceflight and recovered post-return, consistent with circuit-level perturbation and normalisation in the VEN-containing region (Figure 7). This neuroimaging finding is convergent with but methodologically independent of the transcriptomic and proteomic evidence.

**Figure 7.**
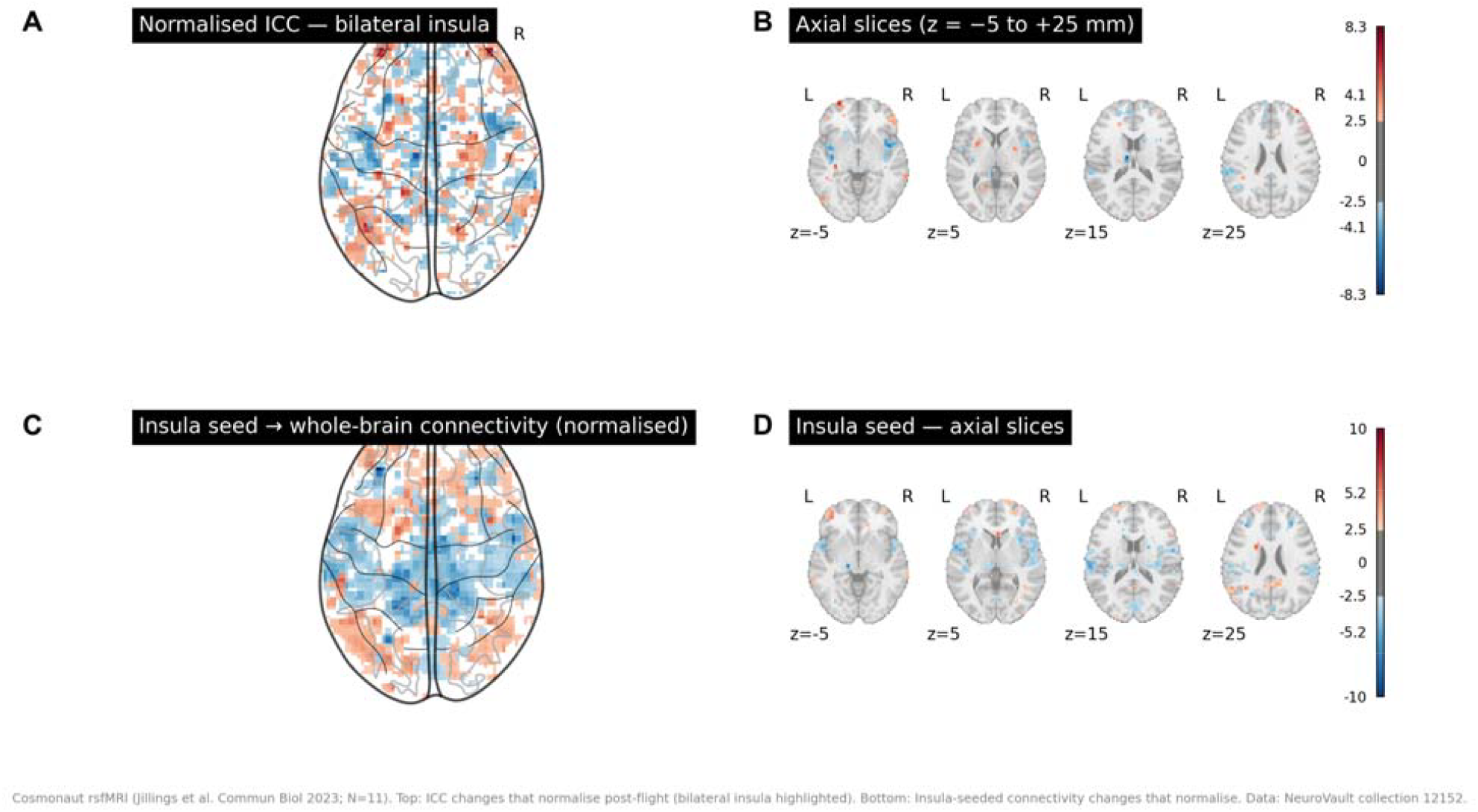
Bilateral frontal insula connectivity is specifically altered by spaceflight and normalises post-return in cosmonauts (Jillings et al., 2023). Resting-state fMRI intrinsic connectivity contrast (ICC) T-maps for the n = 11 cosmonauts who completed a scan at the 8-month follow-up (of the N = 15 scanned before and shortly after flight; NeuroVault:12152 (Jillings et al., 2023)). (A) Normalised ICC glass-brain map (changes recovering to preflight levels by 8-month follow-up), bilateral insula highlighted. (B) Axial slices of the same normalised ICC map, z = −5 to +25 mm. The two largest clusters localise bilaterally to the frontal insula (right: 540 voxels, peak MNI (+48, +2, −6), t = −5.91; left: 428 voxels, peak MNI (−36, −16, 0), t = −5.91). (C) Insula-seeded whole-brain connectivity glass-brain map (same normalised ICC). (D) Axial slices of the insula-seeded map. Colourbar: t-statistic; blue = decreased connectivity post-flight; threshold |t| > 2.5. T-map files reanalysed from NeuroVault:12152 (Jillings et al., 2023) using Python/nilearn by E.K.

### 3.5 Convergence Across Eight Lines of Evidence

Eight convergent but indirect lines of evidence are consistent with the VEN Fatigue Hypothesis: (1) single-subject case observation of duration-dependent ERT pattern; (2) spaceflight-specific rodent myelination upregulation (3.89 SD; p = 0.0001); (3) absence of myelination signal in the radiation-plus-unloading analogue with directional reversal; (4) human cortical organoid Layer V projection gene upregulation (6.17 SD; p = 0.0008); (5) organoid myelination downregulation as the a priori predicted result for OPC-stage models; (6) VEN panel myelination and signalling genes upregulated in human astronaut blood across two independent missions; (7) cross-mission replication of the myelination signal (NASA Twins Study and Inspiration4); (8) bilateral frontal insula connectivity specifically altered and normalised in cosmonauts (Jillings et al., 2023). No single line of evidence is individually sufficient; convergence across species, tissue types, molecular assay types, and neuroimaging modalities constitutes the collective support.

## 4. Discussion

The ERT paradox motivates the VEN Fatigue Hypothesis: performance is stable across 6-month missions, yet declines selectively in the 340-day case observation, while spatial and memory domains remain stable at both time points. The circuit logic rests on two properties unique to VENs. First, their sparse afferent fan-in makes them disproportionately sensitive to input quality: degraded gravity-referenced social cues that would be buffered by thousands of pyramidal afferents instead drive strong VEN responses, generating sustained conduction demand. Second, their metabolically expensive myelination creates a compensatory mechanism with a finite OPC-pool capacity (Monje, 2018). This is distinct from a general white matter demyelination account, which would predict domain-nonspecific slowing; the ERT-selective pattern within the case observation argues specifically against that alternative. Within 6 months, compensatory myelination may sustain ERT speed by accelerating conduction. Beyond a threshold duration, progressive OPC pool depletion may cause fatigue to outpace compensation.

The human evidence is the most direct available. SOMA atlas myelination upregulation (*MOG*: log2FC = +5.82, p = 4.59×10^−27^) replicates across the year-long ISS mission and Inspiration4 (Overbey et al., 2024), spanning different durations, cohorts, and sample types. The blood myelination signal warrants mechanistic comment. *MOG* and *MBP* are myelin structural proteins whose elevation in circulating blood most plausibly reflects active oligodendrocyte myelination or myelin turnover releasing myelin-derived fragments into the bloodstream, a pattern documented during remyelination in white matter injury models (Monje, 2018). Cross-mission replication across the year-long ISS (CPT peripheral blood) and Inspiration4 (PBMC) with distinct sample types, instruments, and analytic pipelines substantially reduces the probability of a single-study artefact. The tissue source of blood *MOG* during spaceflight remains unresolved: peripheral nervous system myelin is an alternative to CNS myelin, and follow-up pairing blood myelin biomarkers with neurofilament light chain would help localise the signal to the brain. Bilateral frontal insula connectivity in cosmonauts was specifically altered post-flight and recovered by 8 months (Jillings et al., 2023), providing neuroimaging evidence of circuit-level perturbation and normalisation in the VEN-containing region which is consistent with a system overdriven during flight and normalising post-return. The cortical organoid Layer V signal (6.17 SD; p = 0.0008) provides human cellular evidence that the neuronal class from which VENs are derived responds specifically to spaceflight conditions. The inverse myelination direction in organoids was correctly predicted a priori: OPC-stage organoids show direct microgravity effects on OPC differentiation, not the activity-driven compensatory myelination expected for mature in vivo VEN circuits. Registration of this prediction before data access (<u>Brain AWG GitHub repository</u>, April 2026) means the organoid result is confirmatory rather than contradictory for the hypothesis.

Competing explanations merit consideration. General cognitive fatigue (Garrett-Bakelman et al., 2019) predicts multi-domain decline, inconsistent with concurrent spatial and memory stability in the case observation. Radiation produces diffuse, domain-nonspecific CNS effects, including long-term neurodegeneration risk (Miller et al., 2026) and broader changes in cognition (Shamsesfandabadi et al., 2024), neither of which would selectively spare other cognitive domains while impairing social emotion recognition specifically. Sleep disruption is pervasive across long-duration spaceflight (Garrett-Bakelman et al., 2019) and frequently invoked as a driver of cognitive decline; however, sleep-deprived performance typically shows domain-general rather than selective slowing, and NASA Twins Study sleep data indicate the most severe disruption occurs in the early mission phase rather than the late inflight phase when ERT speed declined most severely (Garrett-Bakelman et al., 2019). Practice or neuroendocrine effects do not predict a deficit confined to social emotion recognition. The critical internal dissociation is between ISS conditions (microgravity + radiation) and OSD-202 (hindlimb unloading + cobalt-57 radiation): the myelination signal is present under ISS but absent under OSD-202, making the orbital microgravity environment, including vestibular and gravity-referenced social cue disruption, the most parsimonious explanatory variable. This interpretation is independently reinforced by recent findings demonstrating that simulated microgravity directly impairs human midbrain dopaminergic axonal projections (Ali et al., 2026) and drives neurite projection density attrition in the rodent cortex (Pani et al., 2016).

A vestibular disruption account merits separate consideration: chronic otolith dysfunction in microgravity could impair social gaze and gesture processing via cerebellar-insular pathways independently of VEN activity. We propose these accounts are mechanistically complementary rather than competing: disrupted vestibular gravity-referencing constitutes the specific input category that overdrives VEN circuits, given VENs’ established role in integrating bodily-self and social signals in the frontal insula (Allman et al., 2011). Vestibular disruption is thus the upstream cause; VEN fatigue is the downstream consequence. A direct experimental test would compare ERT and non-social spatial tasks under matched vestibular load in a ground-based microgravity analogue.

The most important inferential constraint is the N=1 cognitive dataset, which limits the 340-day ERT finding to a hypothesis-generating case observation. A prospective multi-subject long-duration cohort with concurrent ERT sampling remains the definitive test and the essential next step. Bulk and spatial bulk profiling across all datasets cannot attribute signals specifically to VEN cell bodies rather than co-localised Layer V neurons or oligodendrocytes. The Martins et al. (2026) proteomics dataset (PXD069807) includes both MECP2-deficient and wild-type organoids; MECP2 deficiency substantially alters transcriptional and proteomic profiles, constituting a confound that limits interpretation of the spaceflight proteomics result to the mixed organoid population rather than wild-type alone. Ground-based simulated microgravity applied to iPSC-derived VEN-like neurons via clinostat or rotating wall vessel systems would address this directly, enabling single-cell transcriptomics in the appropriate cell class without requiring spaceflight or post-mortem tissue. Rodent datasets come from species lacking bona fide VENs, meaning extended-duration rodent missions cannot localise signals to VEN cell bodies, though they remain informative for testing the compensatory myelination mechanism through OPC depletion markers (*PDGFRA, NG2*) in frontal cortex from missions exceeding 6 months. The blood myelination signal does not localise the response to the brain, and pairing future measurements with neurofilament light chain would help establish CNS origin. No panel gene is exclusive to VENs, a fundamental constraint shared by any transcriptomic approach to this cell class. Despite these constraints, the duration-threshold prediction is testable within existing ISS mission series without requiring new missions. Sampling ERT performance and blood myelin biomarkers at matched timepoints across 6-month and 12-month rotations in equivalent crews would determine whether performance and biomarker trajectories diverge between durations and whether *MOG* elevation at the inflight second-half predicts within-person ERT speed change. Crewed Mars transit missions of approximately 30 months and planned lunar Gateway rotations exceeding 6 months both cross the proposed duration threshold, and blood myelin monitoring could provide early warning of subclinical ERT decline before mission performance is affected.

## 5. Conclusion

The VEN Fatigue Hypothesis proposes that duration-dependent selective ERT vulnerability in spaceflight arises because VEN circuits, optimised for fast social decisions on sparse input, are overactivated by disrupted gravity-referenced cues, with compensatory myelination sustaining performance within 6 months but failing beyond a duration threshold. Eight convergent but indirect lines of evidence across molecular, cellular, human blood, and human neuroimaging datasets are consistent with this framework. These findings motivate the VEN Fatigue Hypothesis as a target for prospective investigation and provide a circuit-grounded account of duration-dependent cognitive risk relevant to missions beyond the 6-month threshold, including crewed Mars missions of ~30 months.

## Author Contributions

Esila Keskin: Conceptualization, Formal analysis, Investigation, Methodology, Software, Visualization, Writing – original draft, Writing – review and editing.

Margaret Windy McNerney: Data curation, Resources, Supervision, Writing – review and editing.

Nilufar Ali: Data curation, Resources, Supervision, Visualization (figure preparation and BioRender licensing), Writing – review and editing.

## Data and Code Availability

All analysis code, VEN panel definitions, and directional predictions are publicly available at the Brain AWG GitHub repository (https://github.com/OpenScienceDataRepo/Brain_AWG), prospectively deposited prior to data access (April 2026). All datasets are publicly available: GSE239336, OSD-202 (https://doi.org/10.26030/ewfb-7g23), and GSE259421 via NCBI GEO and NASA OSDR; cognitive data via published figures of Garrett-Bakelman et al. (2019) and Dev et al. (2024); proteomics (PXD069807) via PRIDE; blood transcriptomics via the SOMA atlas (Overbey et al., 2024; <u>soma.weill.cornell.edu</u>); cosmonaut rsfMRI via NeuroVault collection 12152 (Jillings et al., 2023).

## Acknowledgements

Research data used in this study were curated by and are available in NASA OSDR (https://science.nasa.gov/biological-physical/data/osdr/). The authors thank the NASA Biological and Physical Sciences (BPS) Division for supporting the NASA Ames Life Sciences Data Archive, NASA GeneLab, and their umbrella project, the NASA Open Science Data Repository. The authors are part of the OSDR Analysis Working Groups (AWGs; https://awg.osdr.space/about), and especially thank the other members of the Brain AWG for the group discussions that informed this work. The AWGs are considered part of NASA Citizen Science.

## Declaration of generative AI and AI-assisted technologies in the manuscript preparation process

During the preparation of this work the author(s) used Anthropic’s Claude (claude.ai) for LaTeX typesetting and Python figure-generation verification. AI assistance was not used to generate scientific content, formulate hypotheses, conduct or interpret statistical analyses, or draw conclusions from data. After using this tool, the author(s) reviewed and edited the content as needed and take full responsibility for the content of the published article. All scientific hypotheses, analysis design and execution, result interpretation, and manuscript writing are the authors’ own.

