## Supplementary file for "The Social Cognition Paradox in Long-Duration Spaceflight: A VEN Fatigue Hypothesis for Duration-Dependent Emotion Recognition Decline"

### S1 Supplementary Methods: Full Data Source Protocols

#### ISS rodent transcriptomics (GSE239336)

ISS frontal cortex data were obtained from GSE239336 (GeoMx Digital Spatial Profiling; ground control vs. spaceflight saline condition; SpaceX CRS-24, 35 days on the ISS; JAXA Life Sciences programme). Frontal cortex tissue was collected post-mission and profiled using GeoMx DSP, a multiplexed spatial transcriptomic platform that quantifies gene expression within defined regions of interest in intact formalin-fixed tissue. Ground control animals were maintained under matched husbandry conditions. The saline-treated spaceflight and ground control arms were compared to isolate spaceflight effects from pharmacological intervention arms. Published log_2_ fold-change values (spaceflight vs. ground) were used directly.

#### Ground-based spaceflight analogue (OSD-202)

OSD-202 was obtained from NASA OSDR (Mao, 2018) (hindlimb unloading combined with cobalt-57 radiation vs. normal loaded control at 1 month). This condition replicates mechanical unloading and radiation exposure without orbital microgravity and without disruption of gravity-referenced social or vestibular cues. Published log_2_ FC values were used directly.

#### Human iPSC-derived cortical organoids (GSE259421)

GSE259421 was obtained from NCBI GEO (Marotta et al., 2024). This dataset comprises bulk RNA-seq from iPSC-derived cortical and dopaminergic organoids maintained 38 days on the ISS or in parallel ground control conditions, with or without iPSC-derived microglia. Organoids at this differentiation stage contain primarily OPCs and immature neurons rather than mature myelinating oligodendrocytes. Cortical organoid samples from two donors yielded nLEO = 9 and ngnd = 10 samples across donor and microglia conditions. Raw count data (61,860 Ensembl-annotated genes) were normalised to CPM and log_2_-transformed (log_2_ (CPM+1)). Ensembl identifiers were mapped to HGNC symbols via MyGene.info (Xin et al., 2016); 45,241 of 61,860 identifiers were successfully mapped (73.2%), yielding 60,799 unique gene symbols. Combined, without-microglia (nLEO = 4, ngnd = 5), and with-microglia (nLEO = 5, ngnd = 5) groups were analysed separately and combined; primary results report the combined group.

#### Human iPSC-derived dopaminergic organoids (GSE259421)

The same GSE259421 dataset contains midbrain-patterned dopaminergic organoids from the same donors and spaceflight conditions. *DRD1* is included in the Social Circuit category because VENs express this receptor as a target for dopaminergic input; VENs do not synthesise dopamine. Sample sizes: without microglia (nLEO = 7, ngnd = 6); with microglia (nLEO = 4, ngnd = 6); combined (nLEO = 11, ngnd = 12). *TH* and *SLC6A3 (DAT)* were tracked as positive controls for dopaminergic identity.

#### iPSC brain organoid proteomics (PXD069807)

Protein-level data from Martins et al. (2026) (PXD069807; PRIDE proteomics repository). Orbitrap Astral LC-MS/MS from WT83 iPSC-derived brain organoids, 30 days ISS vs. ground control, processed with PatternLab V. Identifications filtered at 1% protein FDR. Of the 31 VEN panel proteins, 11 were detected: *CNP* and *ERMN* (Myelination); *ANK3, NEFH, NEFM, NEFL, SNCG* (Fast Signalling); *VDAC1, ATP2B2, SNAP25, SYP* (Metabolic Support). The remaining 20 proteins, including all Layer V transcription factors and Social Circuit receptors, were below the detection threshold; transcription factors are typically expressed at fewer than 1,000 copies per cell, below the single-organoid Orbitrap detection limit.

#### Cognitive data

NASA Twins Study cognitive data were obtained from Figure 10B of Garrett-Bakelman et al. (2019), representing the spaceflight twin’s performance in SD units relative to a 15-astronaut preflight baseline, corrected for the ground control twin. Values for early inflight (months 1-6) and late inflight (months 7-12) were extracted for all 10 Cognition battery subtests via manual digitisation (estimated uncertainty ±0.2 SD). Six-month mission data from Table 2 of Dev et al. (2024) (N = 25 astronauts).

#### Human blood transcriptomics (SOMA atlas)

Obtained from the SOMA atlas (<soma.weill.cornell.edu>; Overbey et al. (2024); Meydan & Mason Lab, Weill Cornell Medicine), aggregating polyA+ blood RNA from the NASA Twins Study (CPT peripheral blood mononuclear cells, year-long ISS) and Inspiration4 (PBMC and cell-free RNA).

#### Cosmonaut resting-state fMRI (NeuroVault:12152)

Resting-state fMRI data from Jillings et al. (2023), publicly available at NeuroVault:12152 (Gorgolewski et al., 2015). Dataset comprises intrinsic connectivity contrast (ICC) T-maps from N = 15 cosmonauts (right-handed males, mean age 46 years) scanned before, shortly after, and 8 months after missions exceeding 3 months, using a 3T GE Discovery MR750 scanner. Six ICC T-maps are available: post vs. pre (all changes); sustained ICC effects (persistent at 8-month follow-up); normalised ICC effects (recover to preflight by 8 months); and seed-ROI analyses for bilateral insula, thalamus, and right angular gyrus. The normalised ICC map (NeuroVault image 773066) was analysed using Python/nilearn to identify cluster coordinates.

### S2 Permutation Test vs Within-Panel t-Test: Complementary Interpretations

The permutation test and within-panel one-sample t-test address complementary questions. The permutation test asks: is the VEN category mean more extreme than expected from any equivalently sized random gene set drawn from the full expressed transcriptome? This tests pathway specificity relative to genome-wide background. The within-panel t-test asks: are individual gene effects within the panel consistently non-zero across panel members? This tests within-category consistency.

For Layer V Projection genes, permutation significance (p = 0.0008) with non-significant within-panel t-test (p = 0.189) reflects signal concentrated in *BCL11B* (+4.12 log_2_FC) and *TBR1* (+2.52) within a five-gene panel. The category mean is extreme relative to the genome-wide background (high permutation significance), but the two large effects are not shared evenly across all five panel members (non-significant t-test). Both statistics are reported in Table S1; permutation results are emphasised because they directly address pathway specificity against the genome-wide null.

### S3 Full Cortical Organoid Results

**Table S1: VEN panel results in human iPSC-derived cortical organoids (GSE259421), ISS vs. ground control.** Significance: ∗ ∗ ∗ *p <* 0*.*001; ∗∗ *p <* 0*.*01; ∗ *p <* 0*.*05; † *p <* 0*.*10 (permutation, two-tailed, *N* = 10*,*000, seed 42).

| **Group** | **Category** | *n* | **Mean FC** | *t* | *pt* | **SD null** | **Perm** *p* |
| --- | --- | --- | --- | --- | --- | --- | --- |
| No microglia | Myelination | 7 | *−*0*.*462 | *−*1*.*69 | 0.141 | *−*2*.*10 | 0*.*050^†^ |
| (*n*LEO = 4, | Fast Signalling | 7 | +0*.*234 | +0*.*66 | 0.535 | +1*.*13 | 0.192 |
| *n*_gnd_ = 5) | Social Circuit | 6 | +0*.*392 | +0*.*61 | 0.571 | +1*.*74 | 0*.*077^†^ |
|  | Layer V Proj. | 5 | +1*.*423 | +1*.*45 | 0.220 | +5*.*72 | 0*.*002^∗∗^ |
|  | Metabolic Supp. | 6 | +0*.*947 | +2*.*37 | 0.064 | +4*.*14 | 0*.*005^∗∗^ |
| With microglia | Myelination | 7 | *−*0*.*443 | *−*1*.*74 | 0.132 | *−*2*.*55 | 0*.*028^∗^ |
| (*n*LEO = 5, | Fast Signalling | 7 | +0*.*018 | +0*.*05 | 0.960 | +0*.*19 | 0.784 |
| *n*_gnd_ = 5) | Social Circuit | 6 | +0*.*242 | +0*.*50 | 0.628 | +1*.*37 | 0.137 |
|  | Layer V Proj. | 5 | +1*.*249 | +1*.*73 | 0.161 | +6*.*27 | *<* 0*.*001^∗∗∗^ |
|  | Metabolic Supp. | 6 | +0*.*530 | +2*.*43 | 0.045 | +2*.*94 | 0*.*017^∗^ |
| Combined | Myelination | 7 | *−*0*.*456 | *−*1*.*78 | 0.125 | *−*2*.*35 | 0*.*035^∗^ |
| (*n*LEO = 9, | Fast Signalling | 7 | +0*.*116 | +0*.*33 | 0.752 | +0*.*66 | 0.387 |
| *n*_gnd_ = 10) | Social Circuit | 6 | +0*.*309 | +0*.*56 | 0.599 | +1*.*56 | 0*.*098^†^ |
|  | Layer V Proj. | 5 | +1*.*347 | +1*.*58 | 0.189 | +6*.*17 | 0*.*0008^∗∗∗^ |
|  | Metabolic Supp. | 6 | +0*.*731 | +2*.*44 | 0.059 | +3*.*60 | 0*.*009^∗∗^ |

Mean FC = mean log_2_FC (ISS−ground); pt = within-panel one-sample t-test; SD null = SDs above genome-wide permutation null.

### S4 Dopaminergic Organoid Results

The same GSE259421 dataset contains midbrain-patterned dopaminergic organoids from identical donors and conditions. Applying the identical VEN panel tests whether signals are cortex-specific.

Layer V Projection genes showed the strongest signal: mean log_2_FC = +1.401 (7.12 SD above null; permutation p = 0.0002), exceeding the cortical organoid signal (6.17 SD; p = 0.0008). Dominant drivers were *BCL11B* (+3.57) and *TBR1* (+2.94), suggesting spaceflight drives a shift toward deep-layer cortical identity programmes not restricted to the cortical lineage. Metabolic Support genes replicated: combined +0.686 (3.63 SD; p = 0.008). Myelination genes were not significantly different from null in dopaminergic organoids (-1.37 SD; p = 0.134), suggesting the cortical myelination signal (-2.35 SD; p = 0.035) reflects an OPC-lineage effect specific to the cortical differentiation protocol. As a positive control, *TH* showed -2.15 log_2_FC (downregulation under spaceflight, consistent with catecholaminergic stress reports); *NOS1* was detected at +0.78 log_2_FC.

**Table S2: VEN panel cross-organoid-type comparison (GSE259421), combined group.** Cortical results from Table S1 shown for comparison.

| **Category** | **Cortical SD** | **Cortical** *p* | **Dopamin. SD** | **Dopamin.** *p* |
| --- | --- | --- | --- | --- |
| Myelination | −2*.*35 | 0*.*035^∗^ | −1*.*37 | 0.134 |
| Fast Signalling | +0*.*66 | 0.387 | +1*.*56 | 0.101 |
| Social Circuit | +1*.*56 | 0*.*098^†^ | +1*.*50 | 0.107 |
| Layer V Proj. | +6*.*17 | 0*.*0008^∗∗∗^ | +7*.*12 | 0*.*0002^∗∗∗^ |
| Metabolic Supp. | +3*.*60 | 0*.*009^∗∗^ | +3*.*63 | 0*.*008^∗∗^ |

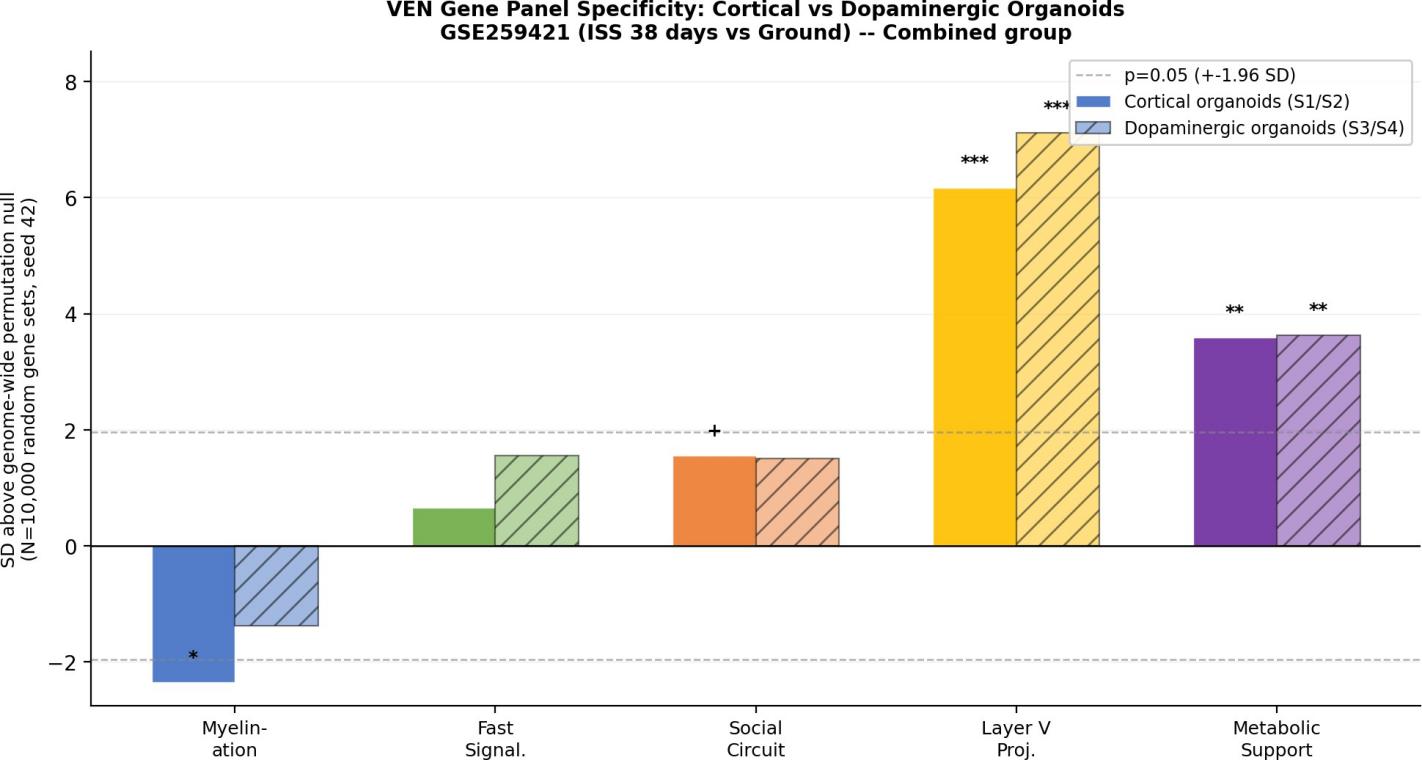

**Figure S1: VEN panel signals replicate across cortical and dopaminergic organoid types (GSE259421).** Standard deviations above the genome-wide permutation null for five VEN gene categories (combined group). Layer V Projection and Metabolic Support signals replicate in both organoid types. Myelination signal present in cortical organoids (−2*.*35 SD, *p* = 0*.*035) is absent in dopaminergic organoids (−1*.*37 SD, p = 0*.*134). ∗ ∗ ∗ p *<* 0*.*001; ∗∗ p *<* 0*.*01; ∗ p *<* 0*.*05; † p *<* 0*.*10 (N = 10*,*000 permutations, seed 42).

### S5 Proteomics: Discordance Between Transcriptomic and Protein-Level Findings

Of the 31 VEN panel proteins, 11 were detected above 1% FDR in PXD069807 (Martins et al., 2026): *CNP* and *ERMN* (Myelination); *ANK3, NEFH, NEFM, NEFL, SNCG* (Fast Signalling); *VDAC1, ATP2B2, SNAP25, SYP* (Metabolic Support). All Layer V transcription factors and all Social Circuit receptors were undetected. Among the 11 detected proteins, all categories showed trends consistent with the genome-wide background downregulation (all-protein mean log_2_FC = −0.42; Myelination detected: −0.60; Fast Signalling: −0.49; Metabolic Support: −0.50). Permutation testing confirmed no category exceeded background (all p > 0.80).

Non-detection of *BCL11B* (+4.12 log_2_CPM in RNA-seq) and *TBR1* (+2.52) is expected: transcription factors are typically expressed at fewer than 1,000 copies per cell, below the single-organoid Orbitrap detection threshold (∼10,000 copies per cell for confident identification). The detection-threshold explanation is biologically plausible but cannot be confirmed without targeted proteomics optimised for low-abundance transcription factors. The protein-level downregulation in detected proteins does not contradict the transcriptomic upregulation, as the two analyses measure different panel subsets at different abundance levels.

### S6 Individual Gene Log_2_FC Table with p-Values

**Table S3: Key individual VEN marker gene** log_2_**FC in human cortical organoids (GSE259421), ISS vs. ground (combined group).**

*p* = Welch two-sample *t*-test (combined group, *n*_LEO_ = 9, *n*_gnd_ = 10; not corrected for multiple testing).

| **Category** | **Gene** | **FC^a^** | **FC^b^** | **FC^c^** | *p* | **Sig.** |
| --- | --- | --- | --- | --- | --- | --- |
| Myelination | *MBP* | −0*.*621 | −1*.*204 | −0*.*941 | 0.0116 | ∗ |
|  | *PLP1* | −1*.*021 | −0*.*702 | −0*.*852 | 0.3580 |  |
|  | *ERMN* | −1*.*765 | −1*.*435 | −1*.*596 | 0.0019 | ∗∗ |
| Fast Signalling | *SCN1A* | +1*.*196 | +0*.*690 | +0*.*929 | *<* 0*.*0001 | ∗ ∗ ∗ |
|  | *NEFM* | +1*.*200 | +0*.*741 | +0*.*955 | 0.0004 | ∗ ∗ ∗ |
| Social Circuit | *CHRM1* | +2*.*935 | +2*.*069 | +2*.*481 | *<* 0*.*0001 | ∗ ∗ ∗ |
|  | *OXTR* | −1*.*741 | −1*.*444 | −1*.*589 | 0.0001 | ∗ ∗ ∗ |
| Layer V Proj. | *BCL11B* | +4*.*497 | +3*.*729 | +4*.*123 | *<* 0*.*0001 | ∗ ∗ ∗ |
|  | *TBR1* | +2*.*929 | +2*.*061 | +2*.*515 | 0.0270 | ∗ |
|  | *FEZF2* | +0*.*448 | +0*.*425 | +0*.*463 | 0.6424 |  |
| Metabolic Supp. | *SLC17A7* | +2*.*825 | +1*.*453 | +2*.*126 | 0.0219 | ∗ |
|  | *NRXN1* | +1*.*102 | +0*.*646 | +0*.*878 | 0.0009 | ∗ ∗ ∗ |
| VEN marker (ind.) | *NOS1* | +0*.*886 | +0*.*234 | +0*.*541 | 0.0018 | ∗∗ |

∗ ∗ ∗ *p <* 0*.*001; ∗∗ *p <* 0*.*01; ∗ *p <* 0*.*05 (Welch two-sample *t*-test, uncorrected). FC^a^ = no microglia (*n*_LEO_ = 4, *n*_gnd_ = 5); FC^b^ = with microglia (*n*_LEO_ = 5, *n*_gnd_ = 5); FC^c^ = combined (*n*_LEO_ = 9, *n*_gnd_ = 10).

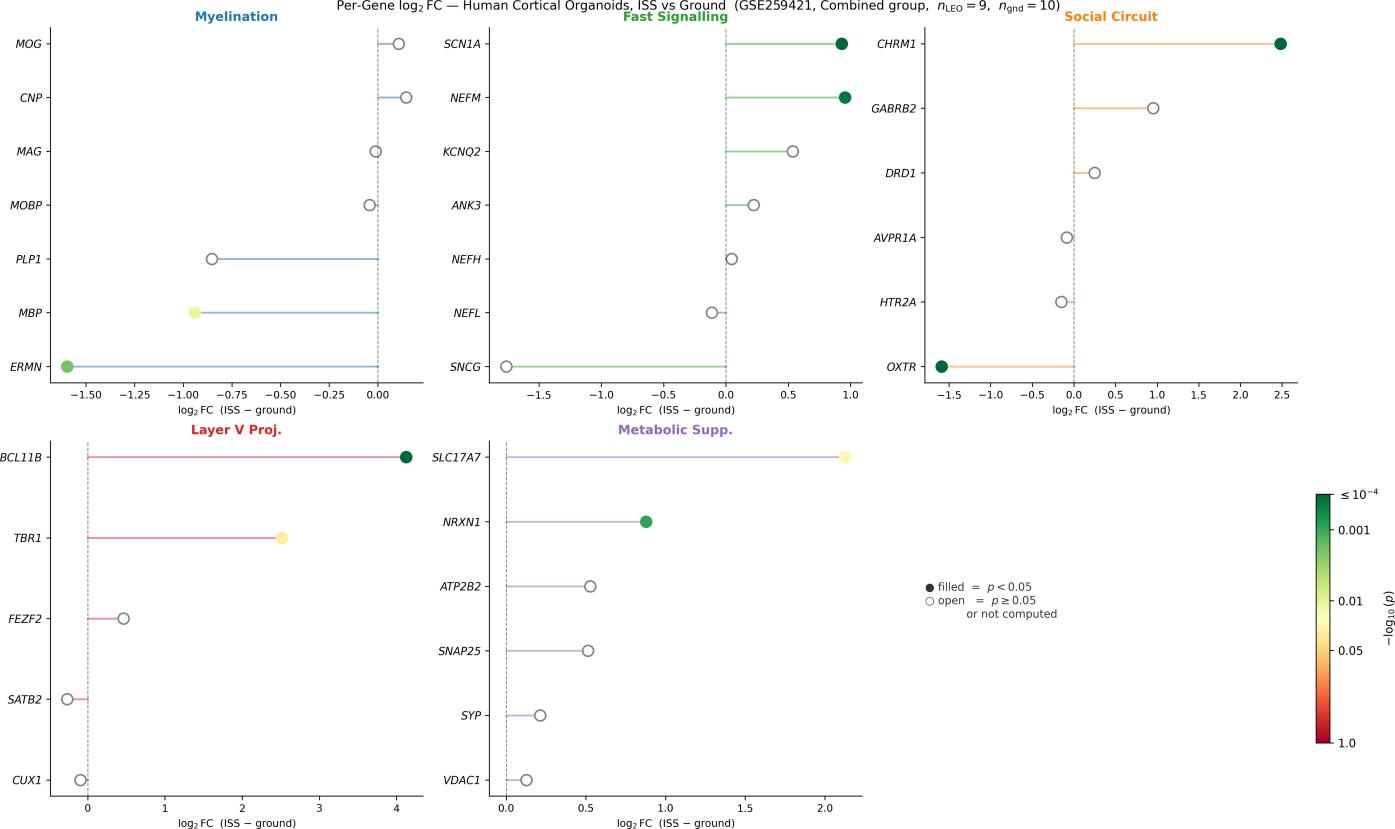

**Figure S2: Per-gene log_2_FC in human iPSC-derived cortical organoids (GSE259421), ISS vs. ground control (Combined group).** Each panel shows all genes in one VEN category. Filled coloured circles: *p <* 0*.*05 (Welch two-sample *t*-test; colour scale = -log_10_(*p*), Red-Yellow-Green). Open grey circles: *p* ≥ 0*.*05 or individual *p*-value not computed for that gene (category-level results reported in Table S1). Values from Combined group (*n*_LEO_ = 9, *n*_gnd_ = 10); all 31 panel genes were detected. Myelination genes are predominantly downregulated (*ERMN, MBP, PLP1*); Layer V Projection genes are dominated by *BCL11B* (+4*.*12) and *TBR1* (+2*.*52).

### S7 Cognitive Domain Specificity (see main text Figure 3)

Early-to-late inflight change scores for all 10 NASA Cognition battery subtests in the single-subject 340-day mission are shown in main text Figure 3 (manually digitised from Garrett-Bakelman et al. (2019) Figure 10B; estimated uncertainty ±0.2 SD). ERT (red) is the most extreme domain by >0.7 SD relative to the within-person non-ERT mean; spatial orientation and visual memory domains remain stable or improve at the same mission phase.

### S8 Cosmonaut rsfMRI: Bilateral Frontal Insula Connectivity (see main text Figure 7)

The ICC maps are shown in main text Figure 7. The bilateral frontal insula and adjacent frontal operculum, the regions housing von Economo neurons (Allman et al., 2011), show significantly reduced connectivity post-flight compared to pre-flight that recovers to preflight levels by the 8-month follow-up. This pattern, perturbation during spaceflight followed by normalisation post-return, is consistent with the VEN Fatigue Hypothesis narrative of circuit-level overactivation during flight and subsequent recovery, though post-flight fMRI does not directly measure inflight dynamics.

The thalamus and right angular gyrus showed sustained connectivity changes that did not normalise by 8 months (NeuroVault images 773076 and 773079). Thalamic sustained changes are consistent with prolonged disruption of vestibular sensory processing; angular gyrus sustained changes are consistent with alterations in multisensory spatial integration. These sustained effects are distinct from the normalised insula signal and may reflect different recovery timescales or mechanisms.
